# Estradiol reverses excitatory synapse loss in a cellular model of neuropsychiatric disorders

**DOI:** 10.1101/455113

**Authors:** Filippo Erli, Alish B. Palmos, Pooja Raval, Jayanta Mukherjee, Katherine J. Sellers, Nicholas J.F. Gatford, Stephen J. Moss, Nicholas J. Brandon, Peter Penzes, Deepak P. Srivastava

**Author notes:** = Authors contributed equally (Filippo Erli, Alish B. Palmos, Pooja Raval). = corresponding author.

## Abstract

Loss of glutamatergic synapses is thought to be a key cellular pathology associated with neuropsychiatric disorders including schizophrenia (SCZ) and major depressive disorder (MDD). Genetic and cellular studies of SCZ and MDD using *in vivo* and *in vitro* systems have supported a key role for dysfunction of excitatory synapses in the pathophysiology of these disorders. Recent clinical studies have demonstrated that the estrogen, 17β-estradiol can ameliorate many of the symptoms experienced by patients. Yet, to date, our understanding of how 17β-estradiol exerted these beneficial effects is limited. In this study, we have tested the hypothesis that 17β-estradiol can restore dendritic spine number in a cellular model that recapitulates the loss of synapses associated with SCZ and MDD. Ectopic expression of wildtype, mutant or shRNA-mediated knockdown of Disrupted in Schizophrenia (DISC1) reduced dendritic spine density in primary cortical neurons. Acute or chronic treatment with 17β-estradiol increased spine density to control levels in neurons with altered DISC1 levels. In addition, 17β-estradiol reduced the extent to which ectopic wildtype and mutant DISC1 aggregated. Furthermore, 17β-estradiol also caused the enrichment of synaptic proteins at synapses and increased the number of dendritic spines containing PSD-95 or that overlapped with the pre-synaptic marker bassoon. Taken together, our data indicates that estrogens can restore lost excitatory synapses caused by altered DISC1 expression, potentially through the trafficking of DISC1 and its interacting partners. These data highlight the possibility that estrogens exert their beneficial effects in SCZ and MDD in part by modulating dendritic spine number.

## Introduction

Glutamatergic synapse dysfunction is thought to be a key cellular hallmark of a number of neuropsychiatric disorders including schizophrenia (SCZ) and major depressive disorder (MDD)^1–4^. In the mammalian forebrain, the majority of excitatory glutamatergic synapses occur on specialized structures known as dendritic spines, which protrude off and decorate the dendrite of pyramidal neurons^4, 5^. Dendritic spines house the post-synaptic density (PSD), where a number of key synaptic proteins involved in synaptic transmission and signaling are located^4, 5^. *Post-mortem* studies have also shown that there are reduced dendritic spine numbers in the brains of patients with SCZ and MDD^2, 6–8^. Consistent with this genetic studies have linked a number of genes encoding for synaptic proteins or for proteins that regulate synapse structure or function with neuropsychiatric disorders including SCZ and MDD^1, 2, 4, 9–11^. Therefore, it has been suggested that regulating synapse protein and dendritic spine number may be a viable therapeutic strategy in the treatment of SCZ and/or MDD^6, 11, 12^.

The *disrupted-in-schizophrenia 1* (*DISC1*) gene has been implicated in the pathophysiology of SCZ and MDD^13–17^. The association of *DISC1* with neuropsychiatric disorders was originally identified through analysis of a large Scottish family where a balanced chromosomal translocation, associated with psychiatric illness^14^. This translocation is thought to lead either to a loss of DISC1 or the formation of a dominant-negative C-terminally truncated DISC1 protein^13, 16^. DISC1 is a scaffold protein that is enriched at synapses where it interacts with a number of different proteins^13, 18^. DISC1 has been described to regulate dendrite spine morphology, number and glutamatergic transmission^13, 15^. While the contribution of *DISC1* to the etiology of SCZ and MDD remains unclear and somewhat controversial^19, 20^, results from animal and cellular models have demonstrated that altering the expression levels of DISC1 protein results in a loss of dendritic spine density^21–24^, a result consistent with that seen in postmortem studies of patients with SCZ or MDD^2, 4, 11^. Truncation of the C-terminal has been used extensively to model DISC1 pathology in cellular and transgenic models. Animal models expressing C-terminal truncated DISC1 constructs have been reported to display reduced spine density *in vivo* as well as *in vitro*^22, 23, 25, 26^.

The neurosteroid, 17β-estradiol, has been shown to be a potent neuromodulator, having positive effects on cognitive processes including learning and memory as well as mood^27, 28^. The effect of 17β-estradiol, the principal biologically active estrogen, on cognition is thought to be driven, in part, by activation of specific signaling pathways resulting in alternations in dendritic spine number and the trafficking of key synaptic proteins (reviewed in 13). Recently, clinical studies have shown that treatment with 17β-estradiol has beneficial effects for patients diagnosed with SCZ or MDD, when given as an adjunct treatment to ongoing antipsychotic or antidepressant therapies^29–34^. However, the molecular and cellular mechanisms by which 17β-estradiol exert these beneficial effects are currently unknown. One possibility is that 17β-estradiol exerts its beneficial effects via the modulation of glutamatergic synapses^27, 35, 36^. However, this has not been tested directly in a cellular model of disease.

In this study, we have tested the hypothesis that 17β-estradiol can restore the number of excitatory synapses in a cellular model that recapitulates the loss of synapses associated with SCZ and MDD. To this end, we have utilized a cellular model of neuropsychiatric disorders, in the form of manipulating the expression levels of DISC1 to reduce dendritic spine density in primary neuronal cultures^21, 23, 37^. Specifically, we have exogenously expressed wildtype rodent DISC1 or a C-terminal truncation mutant of rodent DISC1, which lacks amino acids 598-854, thus mimicking the truncated mutation proposed to be generated by the balance translocation associated with SCZ and MDD, or have used an shRNA approach to knockdown DISC1 expression^13, 21, 38^. Subsequently, we treated cells with 17β-estradiol for 30 minutes (acute) or for 4 days (chronic) to explore whether this neurosteroid could modulate dendritic spine density and synaptic protein expression at synapses in a cellular model, recapitulating aspects of the pathophysiology associated with SCZ and MDD. A number of studies have suggested that the aggregation of DISC1 might be important for psychiatric disease. An original study identified high molecular weight insoluble aggregates of DISC1 in patients specifically diagnosed with major mental illness including SZZ and MDD^39, 40^. Thus, we also investigated whether 17β-estradiol altered mutant or wildtype DISC1 aggregates, and further examined the sub-cellular distribution of endogenous DISC1 and its synaptic interacting proteins following treatment. Our results demonstrate that treatment with 17β-estradiol is sufficient to restore dendritic spine deficits to basal levels, whilst concurrently reducing DISC1 aggregates. Moreover, we go on to show that 17β-estradiol can modulate DISC1 enrichment at synapses, as well as its interacting partner, kalirin-7, a Rac-GEF known to regulate spine density^21^. Critically, we find that chronic treatment with 17β-estradiol increases spine density as well as the number of spines containing PSD-95 and that show overlap with bassoon. Thus, these data suggest that the mechanism by which 17β-estradiol is beneficial in the treatment of SCZ and MDD, may in part be driven by its ability to regulate excitatory glutamatergic synapses.

## Methods

### Reagents

Antibodies used: GFP chicken polyclonal (ab13970; Abcam; 1:10,000); HA mouse monoclonal (clone 16B12; 901503; BioLegend; 1:500); kalirin rabbit polyclonal (07-122; Millipore; 1:500); PSD-95 mouse monoclonal (clone K28/43; 73-028; NeuroMab; 1:1000); PSD-95 guinea pig polyclonal (124014; Synaptic Systems; 1:500); bassoon rabbit monoclonal (6897; Cell Signaling Technologies; 1:200); DISC1 440 rabbit polyclonal (used at 1:250) has been previously described^41, 42^. The antigenic peptide was RTPHPEEEKSPLQVLQEWD, which is identical in human, mouse and rat. DISC1–440 consistently recognizes four major isoforms from rat brain lysates, including two bands at 130 kD and two around 100 kD^41, 42^. 17β-estradiol (E8875) was from Sigma. HA-DISC1WT (wildtype), HA-DISC1ΔCT (mutant) and shRNA against rodent DISC1 constructs were kind gifts from A. Sawa (Johns Hopkins University); generation and validation of these constructs have previously been described^21, 38^. Both wildtype and mutant DISC1 constructs were generated based on rodent sequences; DISC1ΔCT lacks the C-terminal amino acids 598-854, and mimics the breakpoint associated with the balance translocation of DISC1^38, 43^.

### Neuronal culture and transfections

Cortical neuronal cultures, consisting of mixed sexes, were prepared from E18 Sprague-Dawley rat embryos as described previously^44^. Animals were habituated for 3 days before experimental procedures, which were carried out in accordance with the Home Office Animals (Scientific procedures) Act, United Kingdom, 1986. Cells were plated onto 18 mm glass coverslips (No 1.5; 0117580, Marienfeld-Superior GmbH & Co.), coated with poly-D-lysine (0.2mg/ml, Sigma), at a density of 3×10^5^/well equal to 857/mm^2^. Neurons were cultured in feeding media: neurobasal medium (21103049) supplemented with 2% B27 (17504044), 0.5 mM glutamine (25030024) and 1% penicillin:streptomycin (15070063) (all reagents from Life technologies). Neuron cultures were maintained in presence of 200 µM D,L-amino-phosphonovalerate (D,L-APV, ab120004, Abcam) beginning on DIV (days *in vitro*) 4 in order to maintain neuronal health for long-term culturing and to reduce cell death due to excessive Ca^2+^ cytotoxicity via over-active NMDA receptors^44^. We have previously shown that the presence or absence of APV in the culture media does not affect E2’s ability to increase spine linear density^45^. Half media changes were performed twice weekly until desired age (DIV 23-25). The primary cortical neurons were transfected with eGFP at DIV 23 for 2 days, using Lipofectamine 2000 (11668027, Life Technologies)^44^. Briefly, 4-6 µg of plasmid DNA was mixed with Lipofectamine 2000 and incubated for 4-12 hours, before being replaced with fresh feeding media. Transfections were allowed to proceed for 2-5 days after which cells were used for pharmacological treatment or immunocytochemistry (ICC).

### Pharmacological treatments of neuron culture

Acute pharmacological treatments were performed in artificial cerebral spinal fluid (aCSF): (in mM) 125 NaCl, 2.5 KCL, 26.2 NaHCO_3,_ 1 NaH_2_PO_4_, 11 glucose, 5 HEPES, 2.5 CaCl_2,_ 1.25 MgCl_2_, and 0.2 APV). 17β-estradiol was dissolved in DMSO at a concentration of 10 mM, serially diluted to a 10X working concentration in aCSF, and applied directly to neuronal cultures. Final concentration of solvent was < 0.01%; vehicle control was made up of solvent lacking compound, diluted as test compounds. Treatments were allowed to proceed for 30 minutes before being lysed for Western blotting or fixed for ICC.

Chronic pharmacological treatments were performed by adding 17β-estradiol directly into media. Treatment started on the day of transfection and continued ever day until day 4. Cells were fixed 24 hours after the final treatment on day 5. Samples were then processed for ICC.

### Immunocytochemistry (ICC)

Neurons were washed in PBS and then fixed in either 4% formaldehyde/4% sucrose PBS for 10 minutes at room temperature followed by incubation in methanol pre-chilled to −20°C for 10 minutes at 4°C, or only in methanol (−20°C) for 20 minutes at 4°C. Fixed neurons were then permeabilized and blocked simultaneously (2% Normal Goat Serum, 5425S, New England Biolabs and 0.1% Triton X-100) before incubation in primary antibodies overnight and subsequent incubation with secondary antibodies the following day^44^. In the green/magenta colour scheme, co-localization is indicated by white overlap.

### Quantitative Analysis of Spine Morphologies and Immunofluorescence

Confocal images of double-stained neurons were acquired with a Leica SP-5 confocal microscope using a 63x oil-immersion objective (Leica, N.A. 1.4) as a z-series, or with a Zeiss Axio Imager Z1, equipped with an ApoTome using a 63x oil-immersion objective (Carl Zeiss, N.A. 1.4). Two-dimensional maximum projection reconstructions of images were generated and linear density calculated using ImageJ/Fiji (https://imagej.nih.gov/ij/)^44^. Morphometric analysis was performed on spines from two dendrites (secondary or tertiary branches), totaling 100 µm, from each neuron. Linear density and total gray value of each synaptic protein cluster was measured automatically using MetaMorph Software (Molecular Devices)^44^. Cultures directly compared were stained simultaneously and imaged with the same acquisition parameters. For each condition, 10-16 neurons from at least 3 separate experiments were used. Experiments were carried out blind to condition and on sister cultures.

### Biochemistry cell fractionation

Crude synaptosome fractions were prepared from DIV 25 neurons following treatment with 17β-estradiol or vehicle for 30 minutes. Cells were lysed in homogenization buffer (320 mM sucrose; 5 mM Na4P2O7; 1 mM EDTA pH 8; and 10 mM HEPES pH 7.4 + protease inhibitors) and subsequently passed through a 21 gauge needle 15 times. Cell lysate was then centrifuged to remove the nuclear fraction and large cell organelles (P1 fraction) and yield the extranuclear fraction (S1). A portion of the S1 fraction was kept and the remaining was subjected to further fractionation by an additional spin. This yielded the cytosolic (S2) and crude synaptosome (P2) fractions; the P2 fraction was resuspended in homogenization buffer. Sample buffer was added to all samples, which were then denatured for 5 minutes at 95°C and stored at −80°C until used further. All samples were subsequently separated by SDS-PAGE and analyzed by Western Blotting with kalirin, DISC1, PSD-95 and β-actin antibodies. Quantification of bands was performed by measuring the integrated intensity of each band and normalizing to β-actin, for protein loading, using Image J.

### Statistical Analysis

All statistical analysis were performed using Prism (GraphPad Software). Differences in quantitative immunofluorescence, dendritic spine number were identified by Student’s unpaired t-tests, or for comparisons between multiple conditions the main effects and simple effects were probed by one-way or two-way ANOVAs with Tukey correction for multiple comparisons. Error bars represent standard errors of the mean (SEM).

## Results

### Exogenous expression of C-terminal DISC1 mutant causes loss of dendritic spines and aggregates within dendrites

Disruption in glutamatergic transmission and abnormal dendritic spine morphology and number is a cellular phenotype shared across a number of neurodevelopmental and psychiatric disorders^4, 13^. Manipulation of DISC1 function either through expression of wildtype or a C-terminal truncation mutant has previously been shown to induced spine loss *in vitro*^21, 23^. Therefore, we overexpressed either HA-tagged full-length DISC1 (HA-DISC1) or a C-terminal truncated DISC1 mutant (HA-DISC1ΔCT) in DIV 24-26 primary neurons. As expected, neurons expressing HA-DISC1 had a reduced linear spine density compared to control cells (**Figure 1A & B**). Neurons exogenously expressing HA-DISC1ΔCT also displayed reduced spine density compared to control (Spine linear density per 10 µm: one-way ANOVA; *F*(2,46)=21.54, *p*<0.001, η^2^ =0.48; Tukey Post Hoc, ***, p<0.001; n = 12-17 cells per condition from 3 independent cultures; **Supplementary Figure 1A & B**). Recent studies have also indicated that the aggregation of DISC1 may be central to its dominant negative effects and thus, contribute its pathophysiological role^39, 46, 47^. In primary cortical neurons, endogenous DISC1 is localized in punctate structures along dendrites and within dendritic spines, indicating an enrichment at synapses **(Supplementary Figure 2A)**. Consistent with previous reports, HA-DISC1 formed large aggregates within dendrites. Interestingly, HA-DISC1ΔCT also accumulated along dendrites **(Supplementary Figure 2B & C)**. Taken together, these data reveal that both DISC1 and DISC1ΔCT causes a reduction in dendritic spine density and accumulate in dendrites where they appear to form aggregates in primary cortical neurons.

**Figure 1.**
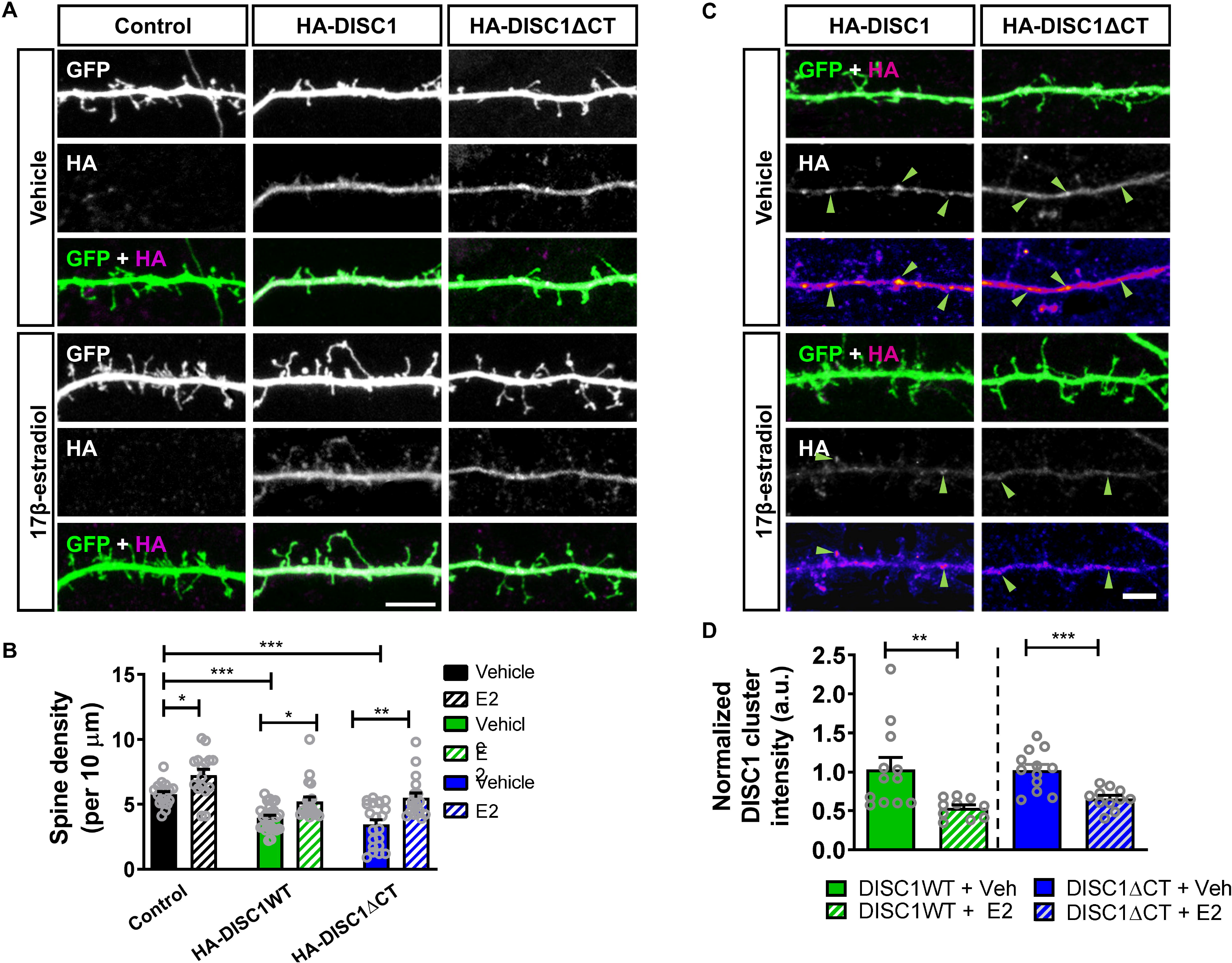
Acute 17β-estradiol (E2) treatment increases dendritic spine linear density in neurons overexpressing HA-DISC1 or HA-DISC1ΔCT and reduces DISC1 aggregation. **(A)** Representative images of cortical neurons (DIV 26) transfected with GFP alone (control), GFP + HA-DISC1 or GFP + HA-DISC1ΔCT and treated with vehicle (Veh) or 17β-estradiol (E2) or not. **(B)** Quantification of spine linear density. Treatment of control cells with 17β-estradiol resulted in an increase in spine number. Overexpression of HA-DISC1 or HA-DISC1ΔCT caused a significant reduction in spine linear density. Treatment with 17β-estradiol increased spine density in HA-DISC1 HA-DISC1ΔCT expressing cells to a level not statistically different to vehicle control cells (dendritic spine linear density/10 µm): control, 5.8±0.21; control + 17β-estradiol, 7.27±0.44; HA-DISC1, 3.9±0.2; HA-DISC1 + 17β-estradiol, 5.22±0.34; HA-DISC1ΔCT, 3.4±0.36; HA-DISC1ΔCT + 17β-estradiol, 5.5±0.37). **(C)** Representative images of DIV 26 cortical neurons expressing either HA-DISC1 or HA-DISC1ΔCT were treated with vehicle or 17β-estradiol for 30 minutes. Pseudo-color scheme is used to highlight regions where DISC1 clustering is occurring. Green arrow heads indicate DISC1 clusters in each condition. **(D)** Quantification of DISC1 clustering; both HA-DISC1 or HA-DISC1ΔCT formed large clusters along dendrites. Treatment with 17β-estradiol reduced DISC1 clustering in neurons expressing either HA-DISC1 or HA-DISC1ΔCT. Scale bar = 5 µm.

### Modulation of Dendritic spine density by 17β-estradiol in neurons expressing wildtype or C-terminal truncated DISC1

Previous studies have demonstrated that 17β-estradiol can increase dendritic spine density in cortical neurons^45, 48^. 17β-estradiol has been reported as having an inverse “U” shaped dose-dependent effect on dendritic spine density^49^. Therefore, we investigated at which dose 17β-estradiol effectively increased dendritic spine density. Cortical neurons expressing GFP were treated either with vehicle (control), or with concentrations of 17β-estradiol: 10 pM, 100 pM, 1 nM, 10 nM or 100 nM for 30 minutes. Analogous to previous studies, we observed an inverse “U” shaped dose-response, where 1 and 10 nM, but not 10, 100 pM or 100 nM, produced a significant increase in spine linear density after 30 minutes of treatment (Spine linear density per 10 µm: one-way ANOVA; *F*(5,97)=17.51, *p*<0.001, η^2^ =0.47; Tukey Post Hoc, **, p < 0.01, ***, p < 0.001; n = 16-21 cells per condition from 4 independent cultures; **Supplementary Figure 3**).

We were next interested in understanding whether 17β-estradiol could modulate dendritic spine density in neurons recapitulating a disease relevant reduction in spine number. Therefore, we next asked whether acute treatment with 17β-estradiol could also restore dendritic spine density in neurons overexpressing HA-DISC1 or HA-DISC1ΔCT. Under control conditions, 17β-estradiol increased spine density compared to vehicle treatment **(Figure 1A, left panel)**. In neurons expressing HA-DISC1, treatment with 17β-estradiol resulted in an increase in the number of spines **(Figure 1A, middle panel)**. Similarly, in neurons ectopically expressing HA-DISC1ΔCT, spine density was again increased following 17β-estradiol treatment **(Figure 1A, right panel)**. Interestingly, spine linear density was similar to untreated control condition in both HA-DISC1 and HA-DISC1ΔCT expressing cells acutely exposed to 17β-estradiol, (Spine linear density per 10 µm: two-way ANOVA; *F*(2,111)=23.73, *p*<0.0001, η_p_^2^ =0.42; Tukey Post Hoc, *, p < 0.05, **, p < 0.01, ***, p < 0.001; n = 16-25 cells per condition from 5 independent cultures; **Figure 1A & B**). Collectively, these data demonstrate that 17β-estradiol is capable of modulating dendritic spine linear density to a level similar to control in neurons expressing either wildtype or mutant DISC1.

### Acute treatment with 17β-estradiol reduces aggregation of wildtype and mutant DISC1

Recent work has suggested that DISC1 forms aggregates under physiological and pathological conditions^34, 41, 42^. The formation of aggregates has been proposed to contributes to DISC1’s cellular function and the cellular pathology associated with the altered expression of this protein^39, 46, 47^. Intriguingly, reducing DISC1 aggregation has been linked with a reduction in cellular pathology^46, 50^. However, whether DISC1 aggregation is linked with the loss of dendritic spine density induced by overexpression, and whether modulating DISC1 aggregation could influence synaptic deficits is not known. As 17β-estradiol could restore aberrant spine density back to a level similar to basal conditions, we tested the hypothesis that it may also reduce the extent to which ectopic DISC1 aggregated. Under control conditions, both HA-DISC1 and HA-DISC1ΔCT could be observed as large clusters within the dendrite (**Figure 1C**). However, after treatment with 17β-estradiol, both HA-DISC1 and HA-DISC1ΔCT clustering was reduced, with smaller and fewer clusters evident within dendrites (HA-DISC1 cluster intensity: Student t-test (corrected for multiple comparisons); t(20)=2.87, p=0.0096, two-tailed, Cohen’s d = 1.28; HA-DISC1ΔCT cluster intensity: Student t-test (corrected for multiple comparisons); t(21)=4.169, p=0.0004, two-tailed, Cohen’s d = 1.96; n = 12-13 cells per condition from 3 independent cultures) **Figure 1C & D**). Taken together, these data indicate that concurrent with the ability to increase dendritic spine density, 17β-estradiol reduces clustering of exogenous wildtype and mutant DISC1.

### Acute 17β-estradiol treatment restores spine loss induced by reduced DISC1 expression

To ensure that the effects observed above were not due to an artifact of overexpressing DISC1 constructs, we employed a second approach to alter DISC1 function. Previous studies have demonstrated that long term shRNA-mediated knockdown of DISC1 results in a loss of dendritic spine linear density in both *in vitro* and *in vivo* systems^21, 37^. Thus, reducing DISC1 expression levels has been suggested to be an approach to induce a cellular phenotype relevant for neurodevelopmental and psychiatric disorders^37^. To this end, we utilized a previously validated shRNA construct for rat DISC1^21, 37, 38^. The DISC1_shRNA construct was able to knockdown exogenous HA-tagged rodent DISC1 in hEK293 cells, confirming it specificity for the protein (Student t-test; t(10)=10.54, p=<0.0001, two-tailed, Cohen’s d = 3.6; **Figure 2A + B)**. We next expressed DISC1_shRNA or control construct in primary cortical neurons for 7 days; this resulted in a significant reduction of spine linear density by ∼35% compared to untreated control **(Figure 2C + D)**, consistent with previous studies^37^. Treatment of control neurons with 10 nM 17β-estradiol increased spine density by ∼40% within 30 minutes **(Figure 2C + D)**. Interestingly, treatment of expressing DISC1_shRNA neurons with 17β-estradiol was sufficient to restore linear spine density to a level similar to control levels (Spine linear density per 10 µm: two-way ANOVA; *F*(1,50)=27.21, *p*<0.0001, η_p_^2^ =0.46; Tukey Post Hoc; * = p < 0.05; *** = p < 0.001; n = 12-15 cells per condition from 4 independent cultures; **Figure 2C + D**). These data demonstrate that 17β-estradiol is capable of modulating spine linear density in neurons with reduced DISC1 levels.

**Figure 2.**
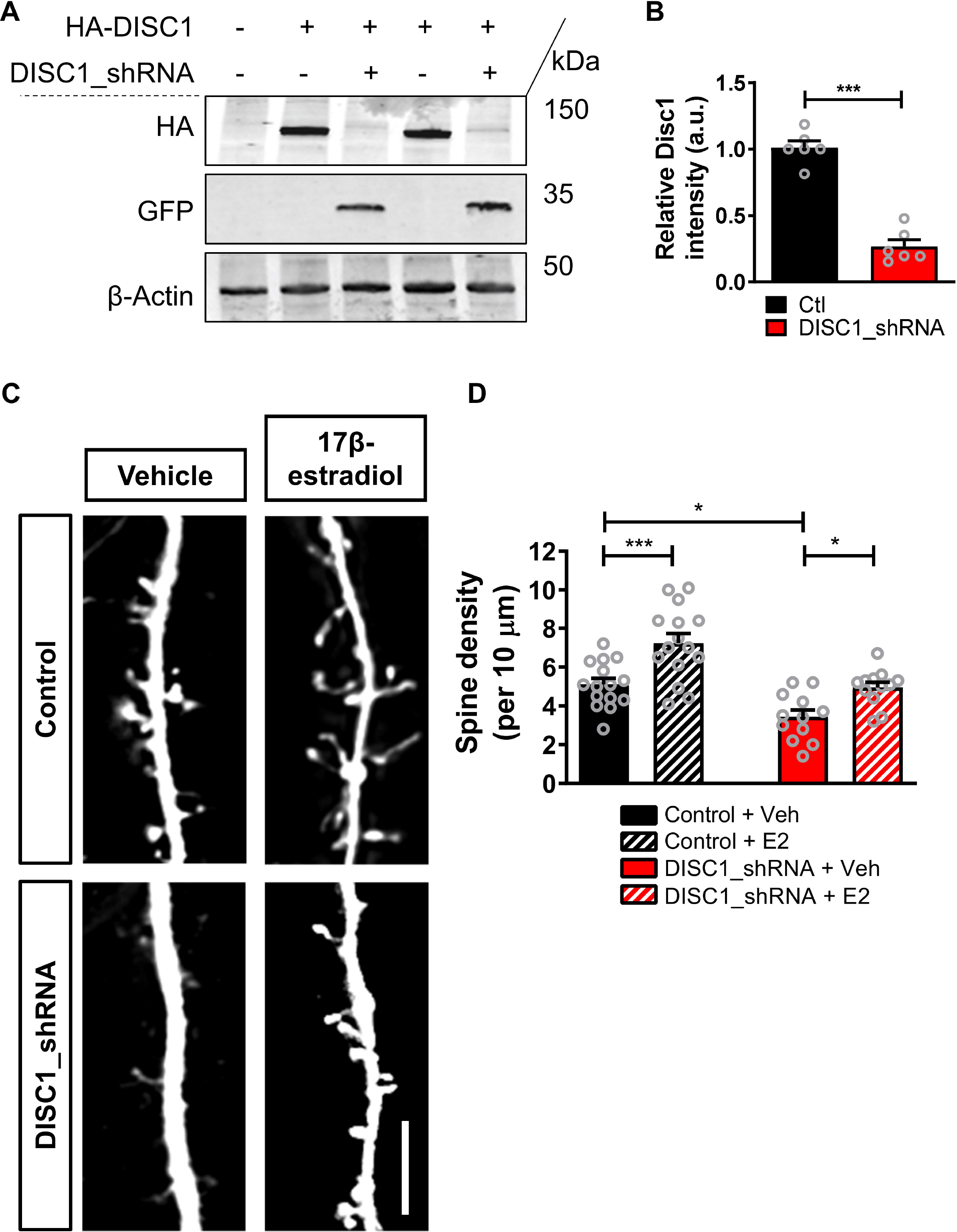
Acute 17β-estradiol (E2) treatment restores dendritic spine density following long term DISC1 knockdown. **(A and B)** Western blot of hEK293 cells expressing HA-DISC1 in presence of control shRNA vector (pGSuper) or DISC1-shRNA **(A)**. Quantification of HA-DISC1 expression **(B)**, normalized to β-actin demonstrates that DISC1_shRNA effectivity knockdown exogenous DISC1 as previously reported. **(C)** Representative images of cortical neurons expressing control shRNA vector (pGSuper) or shRNA against DISC1 (DISC1_shRNA) for 7 days; neurons were treated with vehicle (Veh) or E2 (10 nM) for 30 minutes. **(D)** Quantification of dendritic spine linear density (per 10 µm). As recently reported, expression of DISC1_shRNA for 7 days results in a reduction in spine linear density; as expected 30 minute E2 increases spine density in Control neurons; intriguingly, E2 treatment for 30 min increased spine density in DISC1_shRNA expressing cells to Control neuron levels (dendritic spine linear density (per 10 µm): control, 5.1±0.3; control + 17β-estradiol, 7.25±0.49; DISC1_shRNA, 3.4±0.35; DISC1_shRNA + 17β-estradiol, 4.95±0.27). Scale bar = 5 µm.

### Redistribution of DISC1 and kalirin-7 following acute 17β-estradiol treatment

As our data indicated that 17β-estradiol could modulate the sub-cellular distribution of exogenous wildtype and mutant DISC1, we reasoned that it may also influence the distribution of endogenous DISC1 and its binding partners. Therefore, we evaluated whether acute exposure to 17β-estradiol could alter the sub-cellular localization of the DISC1/PSD-95/kalirin-7 signalosome. In support of this idea we have recently shown that 17β-estradiol causes the rapid redistribution of several synaptic proteins, including PSD-95, from dendrites into spines^48^. Importantly, the ability of DISC1 to regulate spinogenesis is thought to be mediated by its interactions with key synaptic proteins, including PSD-95 and kalirin-7 a Rac-GEF^21^. We first tested this by examining whether levels of DISC1/PSD-95/kalirin-7 were altered following treatment with 17β-estradiol. Western blotting of whole cell lysates of neurons treated with 17β-estradiol for 0 or 30 minutes, revealed no change in overall expression of DISC1 (130 and 100 kDa isoforms), kalirin-7 or PSD-95 (**Figure 3A**; ‘whole cell lysate’). Next, we examined the enrichment of these proteins in crude synaptosomal (P2) and cytosol (S2) fractions. This revealed that 17β-estradiol caused a loss of DISC1 130 and 100 kDa isoforms specifically within the P2 fraction, with a concurrent increase in S2 fraction (**Figure 3A & B**). Conversely, following treatment with 17β-estradiol, the enrichment of both kalirin-7 and PSD-95 in crude synaptosomal fractions (P2) was increased, but decreased in S2 fractions. (DISC1 130 kDa: two-way ANOVA; F(1,8)=24.14, *p*=0.0012, η_p_^2^ =0.75; Bonferroni Post Hoc, *, p < 0.05, **, p < 0.01; n = 3 independent cultures; DISC1 100 kDa: two-way ANOVA; F(1,8)=5.67, *p*=0.0446, η_p_^2^ =0.17; Bonferroni Post Hoc, *, p < 0.05, **, p < 0.01; n = 3 independent cultures; PSD-95: two-way ANOVA; F(1,8)=8.361, *p*=0.0202, η_p_^2^ =0.51; Bonferroni Post Hoc, *, p < 0.05; n = 3 independent cultures; kalirin-7: two-way ANOVA; F(1,8)=9.509, *p*=0.015, η_p_^2^ =0.54; Bonferroni Post Hoc, *, p < 0.05; n = 3 independent cultures; **Figure 3A & B**). To demonstrate that DISC1 was being removed from post-synaptic compartments, we further examined the content of this protein within dendritic spines with or without treatment. As suggested by our biochemical data, assessment of endogenous DISC1 content within spines was significantly reduced following treatment with 17β-estradiol (Student t-test; t(9)=2.97, p=0.0156, two-tailed, Cohen’s d = 1.8; n = 223-337 spines from 5-6 cells per condition from 3 independent cultures); **Figure 3C**). This was further highlighted by line scans analysis across spines, as following 17β-estradiol treatment, the intensity of DISC1 staining within dendritic spines was reduced (**Supplemental Figure 4A**). Thus, these data indicate that 17β-estradiol specifically removes DISC1 from dendritic spines into non-synaptic regions.

**Figure 3.**
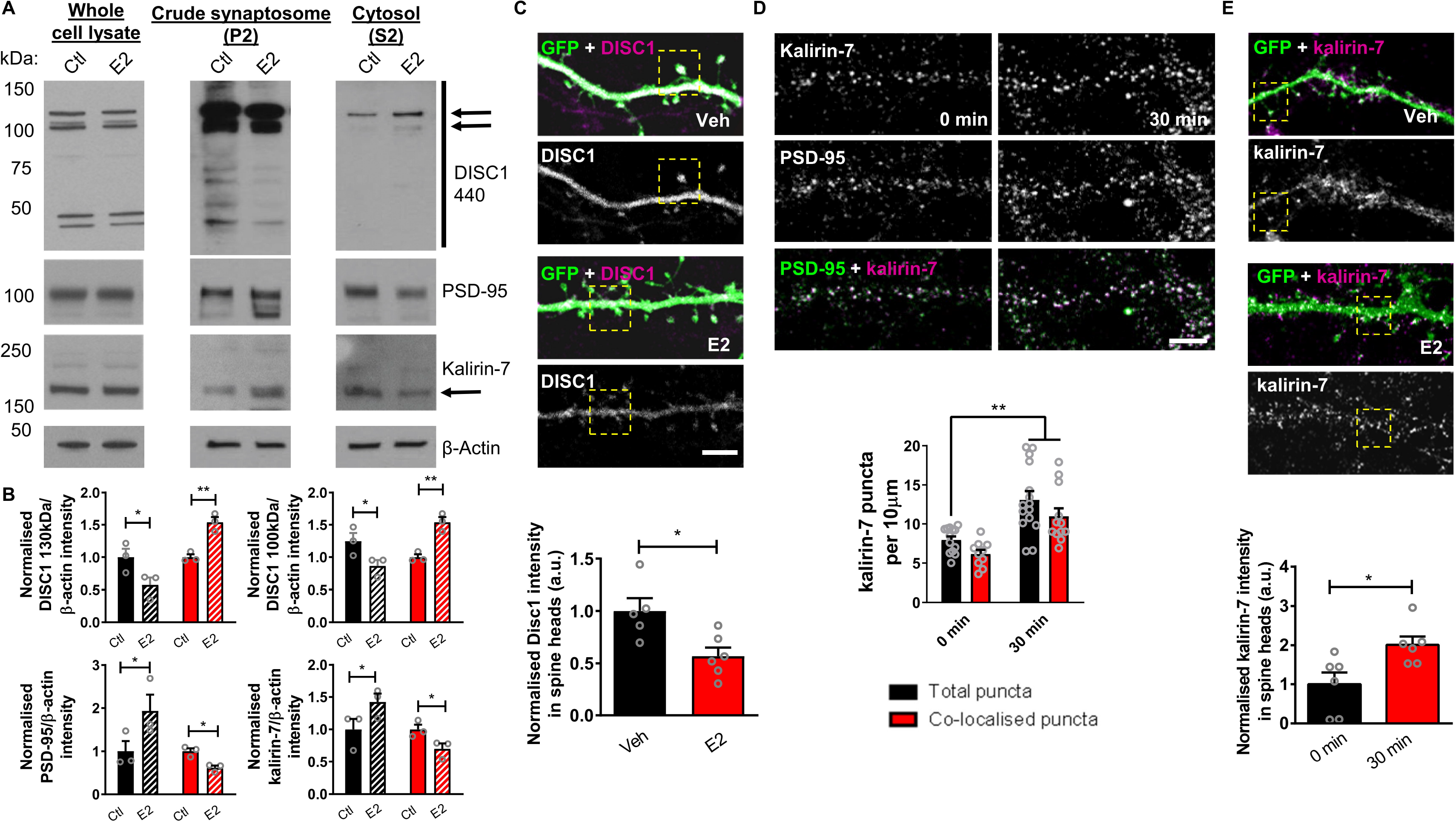
Bi-directional redistribution of DISC1/PSD-95/kalirin-7 signalosome by 17β-estradiol (E2). **(A)** Whole cell lysate, crude synaptosome (P2) and cytosol (S2) fractions of cortical neurons treated with 17β-estradiol (E2) for 0 or 30 minutes analyzed by Western blotting for expression of DISC1, PSD-95 or kalirin-7; β-actin was used as a loading control. Arrows indicate DISC1 and kalirin isoforms measured in analysis. E2 has no effect on overall expression levels of DISC1, kalirin-7 or PSD-95 within 30 minutes as seen in whole cell lysate. Treatment with E2 causes a reduction of DISC1 in crude synaptosome fraction and an increase in cytosolic fraction. E2 treatment resulted in an enrichment of kalirin-7 and PSD-95 within crude synaptosome fraction and concurrent reduction in cytosolic fraction. **(B)** Quantification of DISC1, PSD-95 and kalirin-7 enrichment in crude synaptosome and cytosol fractions. **(C)** GFP-expressing and DISC1 (440) stained cortical neurons treated with E2 for 30 minutes or not. Quantification of DISC1 cluster intensity within spines reveals a reduction in intensity within spines after E2 treatment. **(D)** Cortical neurons (DIV 25) fixed before and after treatment with E2 and immunostaining for kalirin-7 and PSD-95. E2 treatment (30 mins) increases kalirin-7 puncta density (black bars). Assessment of number of kalirin-7/PSD-95 co-localized puncta revealed and increase in kalirin-7 and PSD-95 positive puncta (red bars), indicating that kalirin-7 is being targeted to synapses (*, p < 0.01 (corrected for multiple comparisons) Student t-test; n = 12-15 cells per condition from 3 independent cultures). **(E)** GFP-expressing and kalirin-7 stained cortical neurons treated with E2 for 30 minutes or not. Quantification of kalirin-7 cluster intensity within spines reveals an enrichment within spines following treatment (*, p < 0.05, Student t-test; n = 270-341 spines from 6 cells per condition from 3 independent cultures). Scale bar = 5 µm.

We next sought to determine whether kalirin-7 and PSD-95 were being trafficked to the same location following 17β-estradiol treatment, by assessing the number of kalirin-7 and co-localized PSD-95/kalirin-7 puncta. Consistent with our previous data, we observed an increase in the density of PSD-95 puncta **(Figure 3D)**, which we have previously shown to represent an increase of synaptic sites^48^. Similarly, we found that treatment with 17β-estradiol resulted in an increase in total kalirin-7 puncta (kalirin-7 puncta per 10 µm: two-way ANOVA; *F*(1,46)=4.7, *p*=0.035, η_p_^2^ =0.11; Tukey Post Hoc; ** = p < 0.01; n = 12-15 cells per condition from 3 independent cultures; **Figure 3D**). Under control conditions, 65% of kalirin-7 puncta were positive for PSD-95; a similar level of co-localization was also seen following treatment with 17β-estradiol (**Figure 3D**). Consistent with these data, we found that kalirin-7 intensity within spines increases primarily following 17β-estradiol treatment (**Figure 3E**). The enrichment of kalirin-7 within spines following treatment could also be illustrated using line scan analysis across dendritic spines (**Supplemental Figure 4B**), consequently supporting our biochemical data in demonstrating that a subpopulation of kalirin-7 was being trafficked to synaptic regions following 17β-estradiol treatment. Taken together, these data demonstrate that 17β-estradiol induces a bi-directional trafficking of DISC1, kalirin-7 and PSD-95, with the former protein having a reduced synaptic enrichment, and the latter two proteins increasing in their synaptic localization.

### Chronic repeated treatment with 17β-estradiol reverses spine loss induced by mutant DISC1 expression

We next were interested in understanding whether chronic, repeated exposure to 17β-estradiol could reverse spine loss in our cellular model. To test this, we transfected primary cortical neurons with HA-DISC1ΔCT and GFP or just GFP (control), and simultaneously treated cells with either vehicle or 10 nM 17β-estradiol every day for 4 days. Twenty-four hours after the final treatment (5 days after transfection), cells were fixed, and spine linear density assessed **(Figure 4A)**. In control cells expressing just GFP, repeated exposure to 17β-estradiol produced a 27% increase in spine density. A comparison of vehicle treated control and HA-DISC1ΔCT conditions revealed that mutant DISC expressing cells had 30% fewer spines compared to control neurons. However, repeated exposure to 17β-estradiol reversed this effect and increased spine density in HA-DISC1ΔCT by 30%. Linear density did not differ between control and HA-DISC1ΔCT expressing cells repeatedly exposed to 17β-estradiol (Spine linear density per 10 µm: two-way ANOVA; *F*(1,46)=10.48, *p*=0.0022, η_p_^2^ =0.23; Tukey Post Hoc; * = p < 0.05; n = 12-13 cells per condition from 3 independent cultures; **Figure 4B + C)**. We further examined clustering of exogenous mutant DISC1. This revealed that HA-DISC1ΔCT was less aggregated in cells exposed repeatedly to 17β-estradiol (HA-DISC1ΔCT cluster intensity: Student t-test; t(23)=2.66, p=0.0141, two-tailed, Cohen’s d = 1.07; n = 12-13 cells per condition from 3 independent cultures) **Figure 4B, D & E**). Taken together, these data provide evidence that chronic repeated treatment with 17β-estradiol reverses spine loss induced by mutant DISC1.

**Figure 4.**
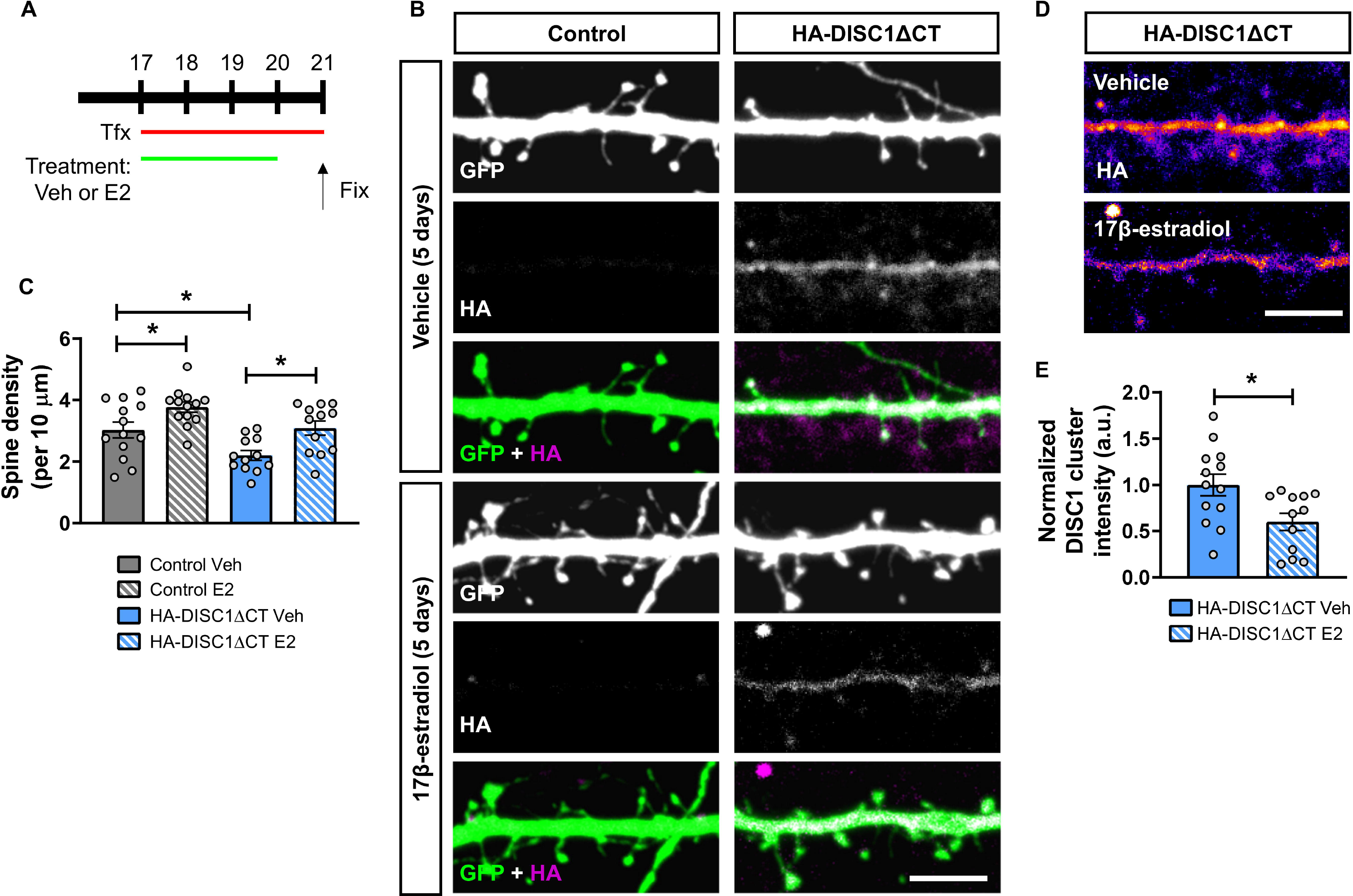
Chronic repeated treatment with 17β-estradiol (E2) rescues DISC1-induced spine deficits and reduces DISC1 aggregation. **(A)** Schematic of chronic repeated 17β-estradiol (E2) treatment of either control or HA-DISC1ΔCT expressing neurons. Cortical neurons were transfected with GFP alone (control) or GFP and HA-DISC1ΔCT at DIV 17. Cells were treated daily with vehicle or 10 nM 17β-estradiol on DIV17-20. Neurons were fixed on DIV21. **(B)** Representative confocal images of DIV25 cortical neurons treated as in **A**. **(C)** Quantification of dendritic spine linear density. Chronic repeated treatment with 17β-estradiol increased in spine density in control cells. Ectopic expression of HA-DISC1ΔCT caused a significant reduction in spine linear density. Chronic repeated treatment with 17β-estradiol increased spine density in HA-DISC1ΔCT expressing cells to a level not statistically different to vehicle control cells. **(D)** Representative images of DIV 25 cortical neurons expressing either GFP alone or HA-DISC1ΔCT were treated with vehicle or 17β-estradiol daily between DIV17 and 24. Pseudo-color scheme is used to highlight regions where DISC1ΔCT clustering is occurring. **(E)** Quantification of DISC1 clustering. Chronic repeated treatment with 17β-estradiol reduced HA-DISC1ΔCT clustering. Scale bar = 5 µm.

One limitation of assessing spine density is that not all spine form synapses. Therefore, we were interested in determining whether chronic, repeated treatment with 17β-estradiol was also altering the number of spines with the potential to form synapses. To this end, we assessed the number of spines that contained PSD-95 or showed overlap with the pre-synaptic and active zone marker, bassoon. Regardless of condition or treatment, we found that approximately 77% of spines were positive for PSD-95 immunoreactive puncta. There was a 26% decrease in the number of spines positive for PSD-95 in HA-DISC1ΔCT expressing cells compared to vehicle treated control neurons **(Figure 5A & B)**. Treatment with 17β-estradiol increased the number of in spines containing PSD-95 by 34% in control cells. In HA-DISC1ΔCT expressing neurons, treatment with 17β-estradiol increased the number of spines containing PSD-95 by 41% compared to vehicle treated HA-DISC1ΔCT expressing cells (Spines positive for PSD-95 density per 10 µm: two-way ANOVA; *F*(1,46)=9.775, *p=0.0031*, η_p_^2^ =0.21; No differences between groups by Tukey Post Hoc; n = 12-13 cells per condition from 3 independent cultures; **Figure 5A & B**). Next we assessed the number of spines that showed overlap for bassoon within the same sample set. Once more, we found no difference in the percentage of spines positive for overlap for bassoon regardless of condition or treatment; approximately 68% of spines overlapped with bassoon across all conditions. Examination of linear density revealed that cells expressing HA-DISC1ΔCT had 20% fewer spines that overlapped with bassoon compared to control neurons **(Figure 5A & C)**. Repeated exposure to 17β-estradiol produced a 41% increase in the number of spines that overlapped with bassoon in compared to both vehicle treated control or HA-DISC1ΔCT expressing cells (Spine overlapping with bassoon density per 10 µm: two-way ANOVA; *F*(1,45)=25.44, *p*<0.0001, η_p_^2^ =0.56 Tukey Post Hoc; * = p < 0.05 ** = p < 0.01; n = 12-13 cells per condition from 3 independent cultures; **Figure 5A & C**). Finally, we assessed the number of spines that were positive for both PSD-95 and that overlapped with bassoon. Across conditions, 60% of spines were positive for PSD-95 and overlapped with bassoon. When we examined the number of spines positive for both synaptic proteins, cells expressing HA-DISC1ΔCT had 30% fewer spines positive for PSD-95 and bassoon compared to vehicle treated control neurons **(Figure 5A & C)**. Neurons repeated exposed to 17β-estradiol had a 50% increase in spines positive for PSD-95 and that overlapped with bassoon compared to vehicle treated control or HA-DISC1ΔCT expressing neurons (Spine positive for PSD-95 and overlapping with bassoon density per 10 µm: two-way ANOVA; *F*(1,46)=23.26, *p<*0.0001, η_p_^2^ =0.51 Tukey Post Hoc; * = p < 0.05 ** = p < 0.01; n = 12-13 cells per condition from 3 independent cultures; **Figure 5A & D**). Collectively, these data provide evidence that chronic repeated exposure to 17β-estradiol is sufficient to increase the number of spines that contain the anatomical hallmarks of being able to form synapses in both control and mutant DISC1 expressing cells. Therefore, 17β-estradiol appears to be able to increase the number of excitatory synapses in a cellular model relevant for psychiatric disorders.

**Figure 5.**
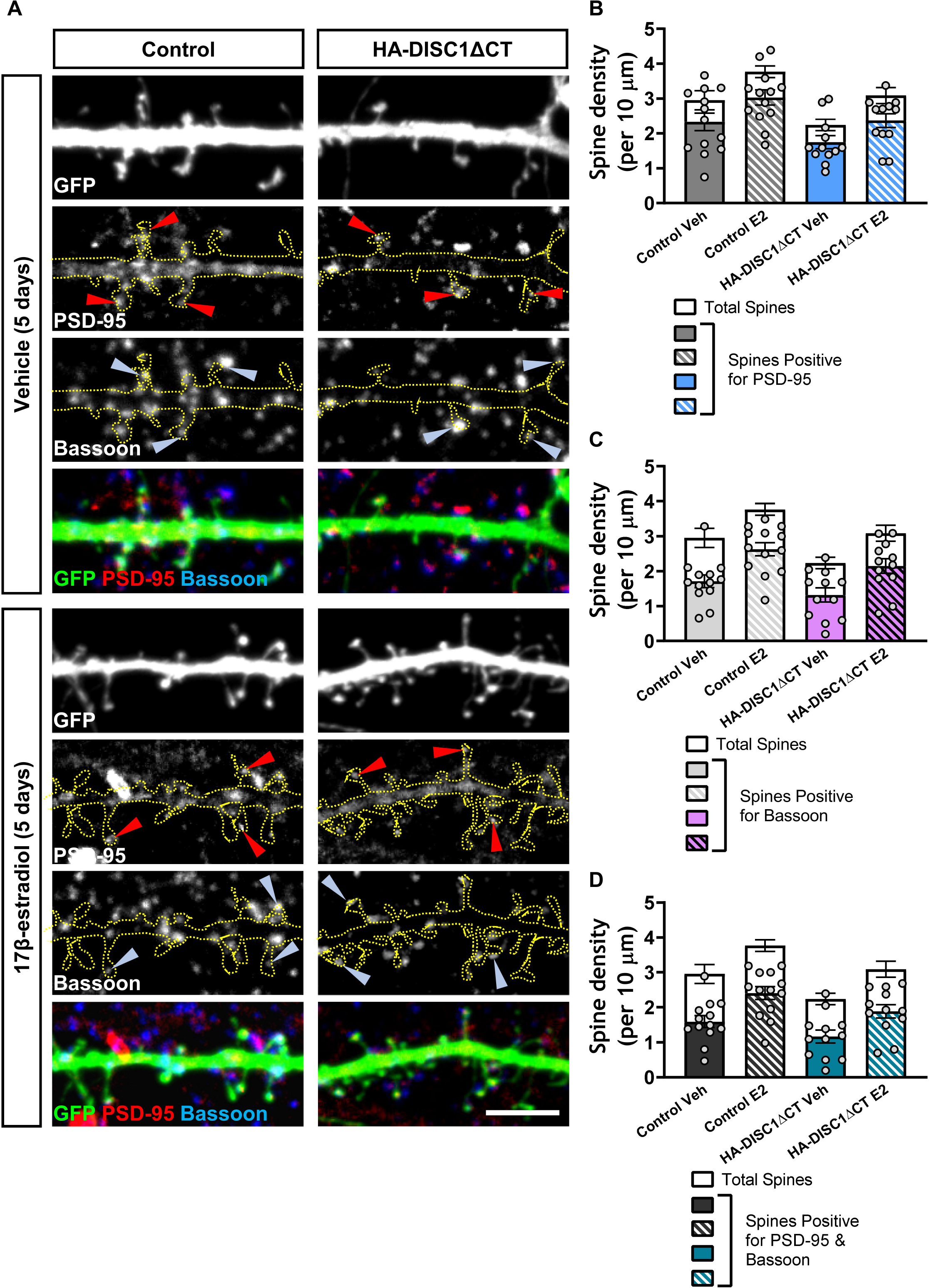
Repeated treatment with 17β-estradiol (E2) increases the number of dendritic spines positive for pre- and post-synaptic markers. **(A)** Representative confocal images of DIV25 cortical neurons triple immunostained for GFP, PSD-95 (post-synaptic protein) and bassoon (pre-synaptic protein). Neurons were treated as outlined in Figure 4A. **(B)** Quantification of linear density of total spine density (open bars) or spines containing PSD-95 (grey and blue solid/hatched bars). Chronic repeated treatment with 17β-estradiol increased the number of spines positive for PSD-95 in control and HA-DISC1ΔCT expressing neurons. **(C)** Quantification of linear density of total spine density (open bars) or spines with overlap for bassoon (light grey and light magenta solid/hatched bars). Chronic repeated treatment with 17β-estradiol increased the number of spines overlapping with bassoon in control and HA-DISC1ΔCT expressing neurons. **(D)** Quantification of linear density of total spine density (open bars) or spines containing PSD-95 and that overlapped with bassoon (dark grey and teal solid/hatched bars). Chronic repeated treatment with 17β-estradiol increased the number of spines positive for PSD-95 and bassoon in control and HA-DISC1ΔCT expressing neurons. Scale bar = 5 µm.

## Discussion

In this study, we have tested the hypothesis that 17β-estradiol can restore excitatory synapses number in a cellular model that recapitulates the loss of synapses associated with SCZ and MDD. We show that in cortical neurons with reduced dendritic spine density produced by manipulating DISC1, acute treatment with 17β-estradiol increased spine density to control levels. Treatment with 17β-estradiol was also able to reduce the extent to which ectopic wildtype and mutant DISC1 aggregated. Furthermore, 17β-estradiol causes a redistribution of DISC1, as well as PSD-95 and kalrin-7, which form a signalsome with DISC1. Finally, we found that chronic, repeated treatment with 17β-estradiol reversed DISC1-induced spine loss in neurons, by increasing the number of spines that also were positive for pre- and post-synaptic protein markers. Taken together, our data indicate that estrogens can restore lost excitatory synapses caused by altered DISC1 expression, potentially through the bi-directional trafficking of DISC1 and its interacting partners.

Recent clinical studies have demonstrated that adjunct treatment with 17β-estradiol and estrogen-based compounds ameliorate positive, negative and cognitive symptoms experienced by patients diagnosed with schizophrenia or depression^29–34^. Recent *in silico-*based studies have also suggested that estrogen-based compounds many confer beneficial effects in SCZ and MDD^51, 52^. This includes identification of selective estrogen receptor modulators as potential repurposable drugs targets for the treatment of MDD^51^. Utilizing an expression-based drug screen on neural progenitor cells derived from induced pluripotent stem cells (iPSCs), generated schizophrenic individuals the estrogenic compound estriol was found to reverse the transcriptional signature associated with SCZ^52^. Together, these clinical and in silico-based studies provide supporting evidence for a potential beneficial role of estrogens in the treatment of MDD and SCZ.

Despite these lines of evidence, the exact molecular and cellular mechanisms by which estrogens exert these beneficial effects are not well understood. The beneficial effects of 17β-estradiol in disorders such as SCZ and MDD have been proposed to occur through the modulation of monoamine transmitter systems such as dopamine and 5-HT, or via an anti-inflammatory mechanism^30, 33, 36, 53^. However, increasing evidence suggests that the beneficial effects of 17β-estradiol may also be in part conferred through the modulation of glutamatergic signaling^27, 35, 36^. This is supported by animal studies which show that 17β-estradiol enhances performance on a number of cognitive tasks, including attention and learning and memory tasks in healthy animals^27, 28, 54^ as well as models of psychosis^30, 55–57^. Importantly, it is the ability of 17β-estradiol to modulate both glutamatergic and GABAergic systems that underlies these enhancing effects^45, 48, 58–61^. This supports the rationale for examining whether 17β-estradiol could regulate excitatory synapses in a disease-relevant cellular model.

It is important to consider the use of DISC1 as the mediator of cellular pathology in our study. DISC1 had emerged as a candidate risk factor for major mental illnesses, including SCZ, autism spectrum disorder, bipolar disorder and MDD^15^. While DISC1 may not directly contribute to the etiology of these disorders^15, 19, 20^, it has been argued that DISC1-based cellular systems are useful in understanding cellular mechanisms relevant for neuropsychiatric disorders^13^, and in exploring potential therapeutic agents^37^. Taking advantage of the ability of DISC1 to induce spine loss, data presented in this study provides evidence that 17β-estradiol is capable of increase glutamatergic synapse number in a disease-relevant cellular model. Collectively, these data indicate that the beneficial effects of estrogens could be driven in part by the regulation of glutamatergic synapses. It should be noted, however, that these results do not negate the possibility that 17β-estradiol also modulates multiple other systems, and this also contributes to its beneficial effects in SCZ and MDD.

It is of note that in our study overexpression of both full length DISC1 or DISC1ΔCT had similar effects to reduce dendritic spine density. DISC1 has been shown to regulate spine and synapse morphology and density through its interacting partners^13, 18, 21, 41^. Therefore, there are multiple mechanisms by which DISC1 could exert its effects on synapses. For example, DISC1 regulates kalirin-7’s GEF activity for Rac1, by modulating the interaction of this protein with PSD-95^21^. Overexpression of DISC1 increases the interaction between kalirin-7 and PSD-95, resulting in a decrease activation of Rac1^21^. DISC1 also interacts and regulates the activity of the synaptic protein TNIK. Regulation of TNIK activity by DISC1 is required for the correct composition of synaptic proteins at synapses and synaptic number^41^. Several studies have shown that expression of mutant DISC1 lacking the C-terminal domain results in a loss of dendritic spines *in vivo* and *in vitro* indicating a critical role for the C-terminal portion of DISC1 in maintenance of spine density^22, 23, 26^. A recent study using a transgenic animal that expresses a C-terminal fragment of DISC1 in an inducible manner, demonstrated that a time-dependent loss of dendritic spine density^25^. As overexpression of the C-terminal fragment is predicted to reduce the ability of DISC1 to interact with binding partners such as NDEL1, this study provides further evidence that the C-terminal and its binding partners are critical for the maintenance and formation of dendritic spines. However, it is also interesting to note that insoluble DISC1, caused by overexpression of DISC1, is less able to bind to interacting partners such as NDEL1^39^, highlighting functional implications of aggregate formation. In our primary neuronal cultures, both DISC1 and DISC1ΔCT constructs formed large clusters, or aggregates, along dendrites, similar to previous reports^39, 46^. Thus, it is possible that aggregation of DISC1 or DISC1ΔCT disrupts the interactome of this protein, and therefore negatively impacts the maintenance of dendritic spines. Interestingly, our data indicates that concurrent with a return of spine density to basal levels, 17β-estradiol also reduces the extent to which DISC1 forms aggregates. While a direct link between DISC1 clustering and spine deficits has not been established, it will be interesting in future studies to ascertain whether these are two separate events, or whether 17β-estradiol’s ability to restore spine density to basal levels is also connected to its ability to reduced DISC1 clustering.

In support of such an idea, we show that 17β-estradiol is capable of altering the distribution of endogenous DISC1 as well. Interestingly, our data indicates 17β-estradiol reduced the synaptic content of endogenous DISC1. This is consistent with previous data indicating that endogenous DISC1 acts to limit spine formation^21^. Activity-dependent signaling reduces the extent to which DISC1 interacts with PSD-95/kalirin-7 which in turns allows for an increase in spine size and number^21, 37^. Consistent with this model, we show that 17β-estradiol caused the bi-directional redistribution of the DISC1/PSD-95/kalirin-7 signalsome. Whereas the presence of endogenous DISC1 at synapses is reduced following 17β-estradiol treatment, both PSD-95 and kalirin-7 are increased at synapses. This is consistent with our previous data showing that PSD-95 is targeted to nascent spines formed by 17β-estradiol^48^. An increased presence of kalirin-7 at synapses would also be consistent with an increase in spine formation. Indeed, we have previously shown that the synaptic localization of kaliirn-7 is important for both spine maintenance and formation^62^. As PSD-95 is required for correct kalrin-7 regulation of dendritic spines, the observed increased co-localized PSD-95/kalirin-7 puncta would indicate that this complex may contribute to the formation of nascent spines. Therefore, in this model, 17β-estradiol acts to release the complex between DISC1 and PSD-95/kalirin-7, in order to allow it to engage with the machinery required for spine formation.

The data presented in this study support a role for estrogenic-modulation of dendritic spines as a potential cellular mechanism by which its beneficial effects in SCZ and MDD may occur. Owing to the link between estrogenic regulation of spines with improvement in cognitive function, especially in learning and memory^45, 60, 61^, as well as emerging data that estrogen-based compounds improve attention and working memory in male and female SCZ patients^29, 31, 33, 36^, it is intriguing to suggest that this is part of the mechanisms that underlies the beneficial actions of estrogens. It is likely that the regulation of spines by 17β-estradiol is only part of the mechanism underlying its beneficial actions. Critically, future studies using genetic models of disease or patient iPSC models should allow for a more faithful recapitulation of both the cellular and complex genetic architecture associated with these disorders. We have shown that neurons derived from iPSC lines are responsive to estrogens^63^, demonstrating that this system could be useful for the study of estrogen-based compounds. Nevertheless, the data in this study support further investigations into estrogenic modulation of glutamatergic signaling and highlights this mechanism as having relevance for the therapeutic potential of estrogens in SCZ and MDD.

In conclusion, our data indicates that estrogens can restore lost excitatory synapses caused by altered DISC1 expression, potentially through the bi-directional trafficking of DISC1 and its interacting partners. These data highlight the possibility that estrogens exert their beneficial effects in SCZ and MDD, in part by modulating excitatory synaptic number.

## Acknowledgements

This work was supported by grants from Medical Research Council, MR/L021064/1, Royal Society UK (Grant RG130856), the Brain and Behavior Foundation (formally National Alliance for Research on Schizophrenia and Depression (NARSAD); Grant No. 25957), awarded to D.P.S.; NIH grants R01MH071316 and R01MH097216 to P.P; K.J.S. was supported by a McGregor Fellowship from the Psychiatric Research Trust (Grant McGregor 97) awarded to D.P.S.; P.R. is funded by a BBSRC-iCASE studentship (BB/M503356/1) awarded to D.P.S. & N.J.B. We thank the Wohl Cellular Imaging Centre for their help with imaging.

## Author Contributions

F.E., A.B.P., J.M., S.J.M., N.J.B., P.P., K.J.S. and D.P.S. designed experiments. F.E., A.B.P., P.R., K.J.S., N.J.F.G. and D.P.S. performed all experiments and subsequent analysis. J.M., S.J.M. and P.P. produced or oversaw production of reagents; J.M., N.J.B., P.P. and D.P.S. wrote the manuscript.

