## Supplemental Information for "Estradiol reverses excitatory synapse loss in a cellular model of neuropsychiatric disorders"

**Supplemental Material**

**Supplemental Figures**

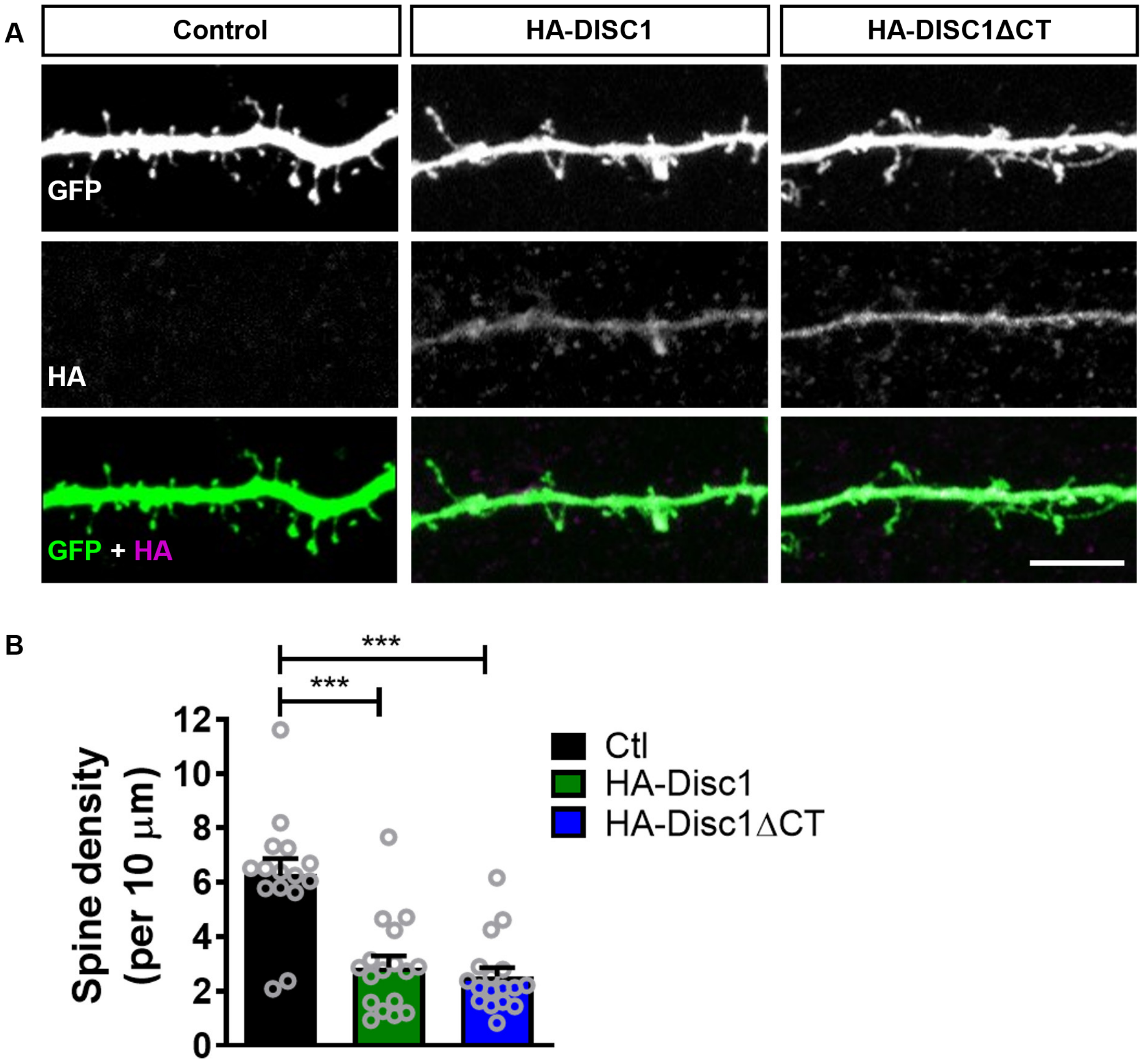


**Supplemental Figure 1. Full length and C-terminal truncated DISC1 induces spine loss in primary cortical neurons.**

**(A)** Representative images of cortical neurons (DIV 26) transfected with GFP alone, GFP + HA-DISC1 or GFP + HA-DISC1ΔCT. **(B)** Quantification of spine linear density. Overexpression of HA-DISC1 causes a reduction of spine density as previously reported; exogenous expression of DISC1ΔCT also causes a reduction in spine linear density (dendritic spine linear density/10 μm): control, 6.3±0.6; HA-DISC1, 2.87±0.42; HA-DISC1ΔCT, 2.54±0.34). Scale bar = 5 μm.

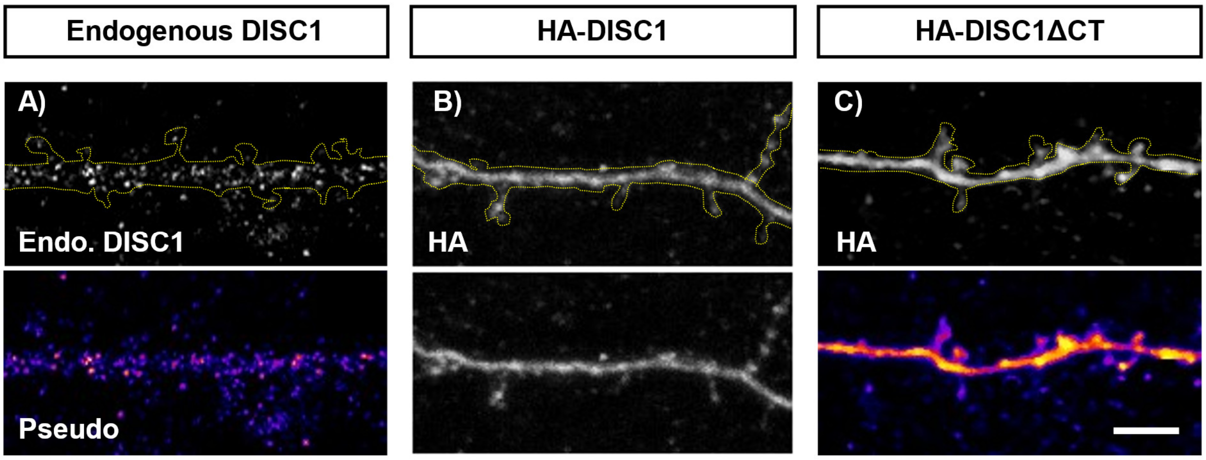


**Supplemental Figure 2. Full length and C-terminal truncated DISC1 aggregates within dendrites of cortical neurons.**

Representative images of cortical neurons (DIV 26) transfected with GFP alone, GFP + HA-DISC1 or GFP + HA-DISC1ΔCT. **(A)** Endogenous DISC1 could be observed in discreet punctate strictures within dendrites and juxtaposed to dendrites. In comparison, both wildtype and ΔCT forms of DISC1 were found to aggregate within dendrites. Scale bar = 5 μm.

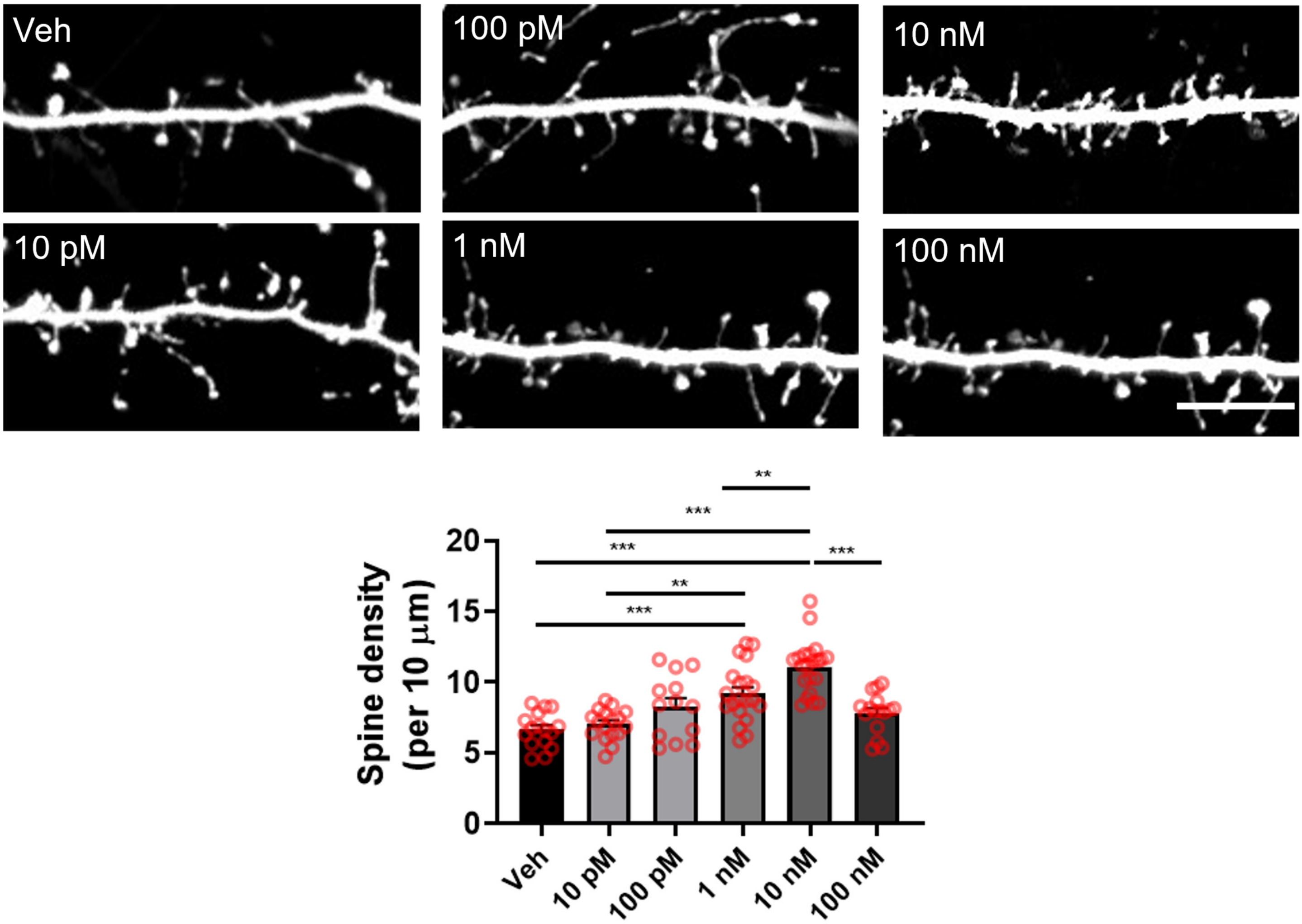


**Supplemental Figure 3. 17β-estradiol increases dendritic spine density in a dose-response manner.** Primary cortical neuron expressing GFP were treated with 17β-estradiol at range of concentrations for 30 minutes. Following fixation and immunocytochemistry, dendritic spine linear density was assessed. Treatment with 1 or 10 nM 17β-estradiol increased spine linear density significantly; 10 or 100 pM as well as 100 nM 17β-estradiol did not alter spine density significantly compared to vehicle treated control neurons. Scale bar = 5 µm.

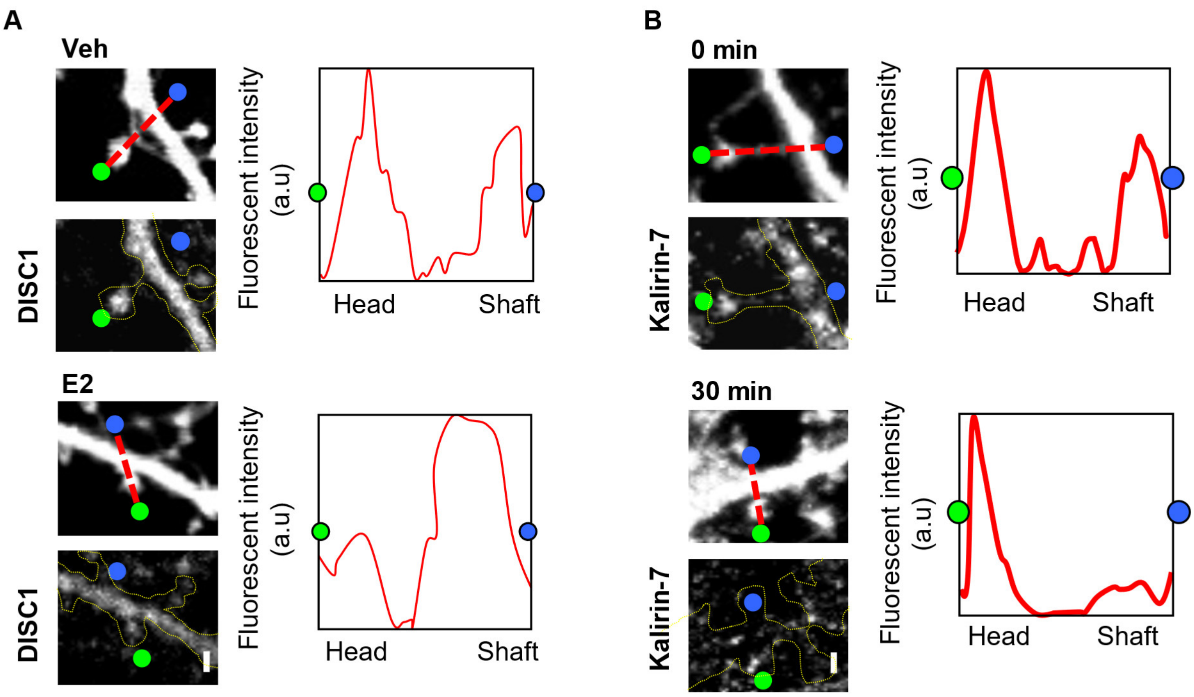


**Supplemental Figure 4. Estradiol induced trafficking of DISC1 and kalirin-7. (A)** High magnification of areas highlighted by yellow dashed boxes in **Figure 3C**; line scan of DISC1 staining within spine heads further demonstrates a redistribution of DISC1 from spines to dendritic shaft after E2 treatment. **(B)** High magnification of areas highlighted by yellow dashed boxes in **Figure 3E**; line scan of kalirin-7 distribution within spine heads reveals an enrichment of kalirin-7 within spine heads after E2 treatment. Scale bar = 1 µm.
